# Metabolic and transcriptomic profiles of glioblastoma invasion revealed by comparisons between patients and corresponding orthotopic xenografts in mice

**DOI:** 10.1101/2021.06.29.450408

**Authors:** Cristina Cudalbu, Pierre Bady, Marta Lai, Lijing Xin, Olga Gusyatiner, Marie-France Hamou, Mario Lepore, Jean Philippe Brouland, Roy T. Daniel, Andreas F. Hottinger, Monika E. Hegi

**Affiliations:** CIBM Center for Biomedical Imaging, Switzerland; Animal Imaging and Technology, Ecole Polytechnique Fédérale de Lausanne (EPFL), Lausanne, Switzerland; Swiss Institute of Bioinformatics (SIB), Lausanne, Switzerland; Neuroscience Research Center, Lausanne University Hospital (CHUV) and University of Lausanne (UNIL), Lausanne, Switzerland; Laboratory for Functional and Metabolic Imaging, Ecole Polytechnique Fédérale de Lausanne (EPFL), Lausanne, Switzerland; Neurosurgery, Lausanne University Hospital (CHUV) and University of Lausanne (UNIL), Lausanne, Switzerland; Institute of Pathology, Lausanne University Hospital (CHUV) and University of Lausanne (UNIL), Lausanne, Switzerland; Neurology, Lausanne University Hospital (CHUV) and University of Lausanne (UNIL), Lausanne, Switzerland; Oncology, Lausanne University Hospital (CHUV) and University of Lausanne (UNIL), Lausanne, Switzerland; Swiss Cancer Center Léman (SCCL), Lausanne, Switzerland

**Author notes:** **Corresponding author**: Monika E. Hegi, PhD, Clinical neuroscience research Center and Neurosurgery, Lausanne University Hopital and University of Lausanne, Chemin des Boveresses 155, CLE-C306, 1066 Epalinges, Switzerland). Equal contribution (Cristina Cudalbu, Pierre Bady). shared senior authorship (Andreas F. Hottinger, Monika E. Hegi).

**Keywords:** glioblastoma, invasion, high resolution ^1^H-MRS, transcriptome, patient-derived orthotopic xenografts (PDOX), tumor host interaction

## Abstract

The invasive behavior of glioblastoma, the most aggressive primary brain tumor, is considered highly relevant for tumor recurrence. However, the invasion zone is difficult to visualize by Magnetic Resonance Imaging (MRI) and is protected by the blood brain barrier, posing a particular challenge for treatment. We report biological features of invasive growth accompanying tumor progression and invasion based on associated metabolic and transcriptomic changes observed in patient derived orthotopic xenografts (PDOX) in the mouse and the corresponding patients’ tumors. The evolution of metabolic changes, followed *in vivo* longitudinally by ^1^H-Magnetic Resonance Spectroscopy (MRS) at ultra-high field, reflected growth and the invasive properties of the human glioblastoma transplanted into the brains of mice (PDOX). Comparison of MRS derived metabolite signatures, reflecting temporal changes of tumor development and invasion in PDOX, revealed high similarity to spatial metabolite signatures of combined multi-voxel MRS analyses sampled across different areas of the patients’ tumors. Pathway analyses of the transcriptome associated with the metabolite profiles of the PDOX, identified molecular signatures of invasion, comprising extracellular matrix degradation and reorganization, growth factor binding, and vascular remodeling. Specific analysis of expression signatures from the invaded mouse brain, revealed extent of invasion dependent induction of immune response, recapitulating respective signatures observed in glioblastoma. Integrating metabolic profiles and gene expression of highly invasive PDOX provided insights into progression and invasion associated mechanisms of extracellular matrix remodeling that is essential for cell-cell communication and regulation of cellular processes. Structural changes and biochemical properties of the extracellular matrix are of importance for the biological behavior of tumors and may be druggable. Ultra-high field MRS reveals to be suitable for *in vivo* monitoring of progression in the non-enhancing infiltration zone of glioblastoma.

## Introduction

Management of patients suffering from glioblastoma (GBM, WHO grade IV), the most common and most malignant primary brain tumor in adults remains a challenge. Even with the latest treatment options, the median survival remains below two years [1]. This poor outcome has been attributed to the hallmarks of GBM that comprise a plethora of altered pathways, intra-tumoral genetic and metabolic heterogeneity, the interaction with the tumor microenvironment, and characteristic properties of tumor stem like cells [2–4]. These confer properties relevant for the invasive behavior and cell plasticity that render GBM particularly resistant to treatments [5, 6]. The diffuse infiltration of gliomas into the surrounding brain precludes complete resection and gives rise to tumor recurrence even in macroscopically fully resected tumors.

Importantly, the extent of invasion is not visible using conventional T1 and T2-weighted Magnetic Resonance Imaging (MRI). Hence, it is difficult to target treatment to this “invisible” part that has a distinct microenvironment and is protected by an intact blood brain barrier (BBB).

Patient derived orthotopic xenograft (PDOX) models, implanting glioma stem-like cells or glioma derived spheres into the brain of immune-compromised mice, are frequently used to study tumor relevant molecular mechanisms and to evaluate novel therapies. The implanted cells give rise to tumors that recapitulate many relevant characteristics of the parental tumors, including invasiveness [7–10]. Glioma cells are metabolically reprogrammed in order to support proliferation, migration and invasion, and to survive in a hostile environment lacking oxygen and nutrients [11–14]. *In vivo* detection of altered metabolic profiles in GBM, which can be achieved with ultra-high field ^1^H-MR spectroscopy (^1^H-MRS) yielding highly resolved spectra provides an opportunity to gain insights into the mechanisms of tumor progression [15–17]. This allows quantification of molecules involved in energy metabolism, myelination, neurotransmission, antioxidation, and osmoregulation, as determined in rodent xenograft models of glioma [18–21]. The non-invasiveness of the technique allows longitudinal measurements of the metabolism during glioma development in the natural environment, while simultaneously probing the metabolism of the surrounding “non-tumoral” brain [21].

In this study we characterize molecular features associated with highly invasive growth of GBM that may serve as markers of early relapse or response to therapy, and may allow to shed light on underlying biological processes of tumor-host interaction. To this end, we investigated GBM in patients, paired with follow-up of their respective PDOX upon transplantation into the brains of immunocompromised mice. Analyses and comparisons included radiologic evaluation using ultra-high field MRI and ^1^H-MRS of the patients (7 Tesla) and longitudinal follow-up of the respective developing PDOX (14.1 Tesla). Subsequently, we associated the metabolic profiles obtained by ^1^H-MRS with the corresponding transcriptomes to gain insights into molecular mechanisms of tumor-host interaction of GBM invasion. Integrating metabolite profiles with the associated transcriptome allowed novel insights into the biological interactions, while distinguishing contributions originating from the human tumors and the invaded mouse brain, respectively.

## Results

To determine metabolite profiles across distinct parts of unresected GBM, patients with suspected GBM were enrolled into the study and underwent scans at 7 Tesla prior to planned surgery. Highly resolved spectra were obtained using single-voxel and multi-voxel MRS (MVS) (Table 1). Depending on the patient’s clinical condition and tumor location, not all analyses were possible. For nine patients enrolled, fresh tumor tissue became available at surgery for orthotopic transplantation into mice. Six of the patients’ tumors were subsequently diagnosed as GBM, and three as astrocytoma grade III, whereof one was diagnosed as IDHmt, 1p/19q non-codeleted. The patient baseline description is summarized in Table 1.

**Table 1.** Baseline information of patients and PDOX

| Pat ID | Sex | Age | 7 Tesla MVS | Diagn | IDHmt R132H | EGFR (IHC) Hirsch Score<sup>b</sup> |  | No mice inj. | No events | PDOX latency (median) |  | RNA-seq |  |
| --- | --- | --- | --- | --- | --- | --- | --- | --- | --- | --- | --- | --- | --- |
|  |  |  |  |  |  | hu Tu | Xeno |  |  | Days | 95 LCL<sup>f</sup> | hu Tu | Xeno |
| P1 | M | 68 | 1 | GBM | no | 400 | 400 | 6 | 6 | 107.5 | 97 | 1 | 6 |
| P4 | M | 55 |  | GBM | no | <100<sup>c</sup> | 300 | 3 | 3 | 141.0 | 138 | 1 | nd |
| P6 | F | 76 |  | AIII | no | 0 | 0 | 4 | 4 | 71.5 | 63 | 1 | nd |
| P7 | M | 67 |  | AIII | no | 400 | 400 | 5 | 2<sup>d</sup> | 75 | 75 | 1 | 5 |
| P8 | M | 73 |  | GBM | no | 300 | 400 | 5 | 3<sup>d</sup> | 142.5 | 67 | 1 | nd |
| P9 | M | 49 |  | GBM | no | 0 | 100 | 5 | 2<sup>d</sup> | 134.5 | 125 | 1 | nd |
| P10 | F | 38 |  | AIII<br>IDHmt,<br>1p/19q<br>non-codel | yes | 150 | na | 4 | 0<sup>e</sup> | na |  | nd |  |
| P12 | M | 37 | 1<sup>a</sup> | GBM | no | 400 | 0 | 6 | 5 | 53.0 | 34 | 1 | 5 |
| P14 | M | 65 | 1 | GBM | no | 0 | nd | 5 | 3 | 172.0 | 169 | 1 | 3 |

<sup>a</sup>MVS, 6 months after tumor resection, during 3<sup>rd</sup> temozolomide maintenance cycle; <sup>b</sup>Hirsch score [22], range 0 to 400; <sup>c</sup>patchy EGFR expression (0 to 400); <sup>d</sup>censored mice died of non-tumor related experimental complications; <sup>e</sup>no tumor development, follow-up up to 241 days; <sup>f</sup>lower 95% confidence latency
Abbrev: EGFR, epithelial growth factor receptor; hu Tu, human tumor; MVS, multi-voxel spectroscopy; na, not applicable; nd, not done; Xeno, tumor Xenograft in the mouse brain.

### Metabolic profiles of tumor development and invasion in PDOX

After stereotactic transplantation of the patient derived GBM cells into the striatum of 4 to 6 immunocompromised mice, the tumor development was followed longitudinally by MRI and ^1^H-MRS at ultra-high magnetic field (14.1T). First scans were performed 4 to 6 weeks after transplantation, and measurements were repeated thereafter at one to two week intervals, placing the voxel in the presumed injection site and symmetrically in the contralateral side of the brain. The acquired highly resolved spectra, allowed measurements of 21 metabolites (Fig. 1A, B; Supplementary Table 1). Temporal changes of the metabolite profiles monitored on the injected and contralateral sides, were in general more sensitive for early detection of tumor growth than standard clinical MRI acquisitions performed in parallel. Eventually tumor development was revealed for all patients’ samples, except for the IDH-mutant astrocytoma grade III (patient 10, P10). The growth kinetics of the tumors and the spread to the contralateral side was reflected in the evolution of the metabolite profiles over time (Fig. 1C-D). In contrast, the metabolite profiles of the mice transplanted with cells from P10 remained at base line, over the observation time of up to 250 days, in line with the lack of tumor development (Fig. 1D). The metabolite profiles from the injected and contralateral side are visualized for each mouse and measurement in function of time from injection, stratified by patient (Fig. 1D). These metabolite profiles are represented by their coordinates on the first axis of the STATIS compromise analysis (similar to PCA) including all measurements. The corresponding representation of the metabolites on the first axis is depicted in Fig. 1E. Growth and invasion were associated with decreasing neuronal metabolism (N-acetyl aspartate, NAA; glutamate, Glu; and gamma aminobutyric acid, GABA) and increase in metabolite markers specific for high cellular turnover such as choline compounds (total choline, tCho; glycerophosphocholine, GPC) and myo-inositol (Ins) as visualized by their coordinates on the first axis of STATIS (Fig. 1E). High values on this axis correspond to high “tumoral properties”, while low values are associated with more normal features, resembling the profiles of normal brain. Accordingly, small or no changes in the metabolite slopes registered in the contralateral side of xenografts reflected absence or lower (e.g. P12, P14) infiltrative capacity as confirmed by histology (Fig. 1C-D). In line, no changes from baseline (“normal”) were observed on either side of the brain in mice orthotopically injected with cells from P10 that did not form any histologically detectable tumors over the observation period.

**Fig. 1.**
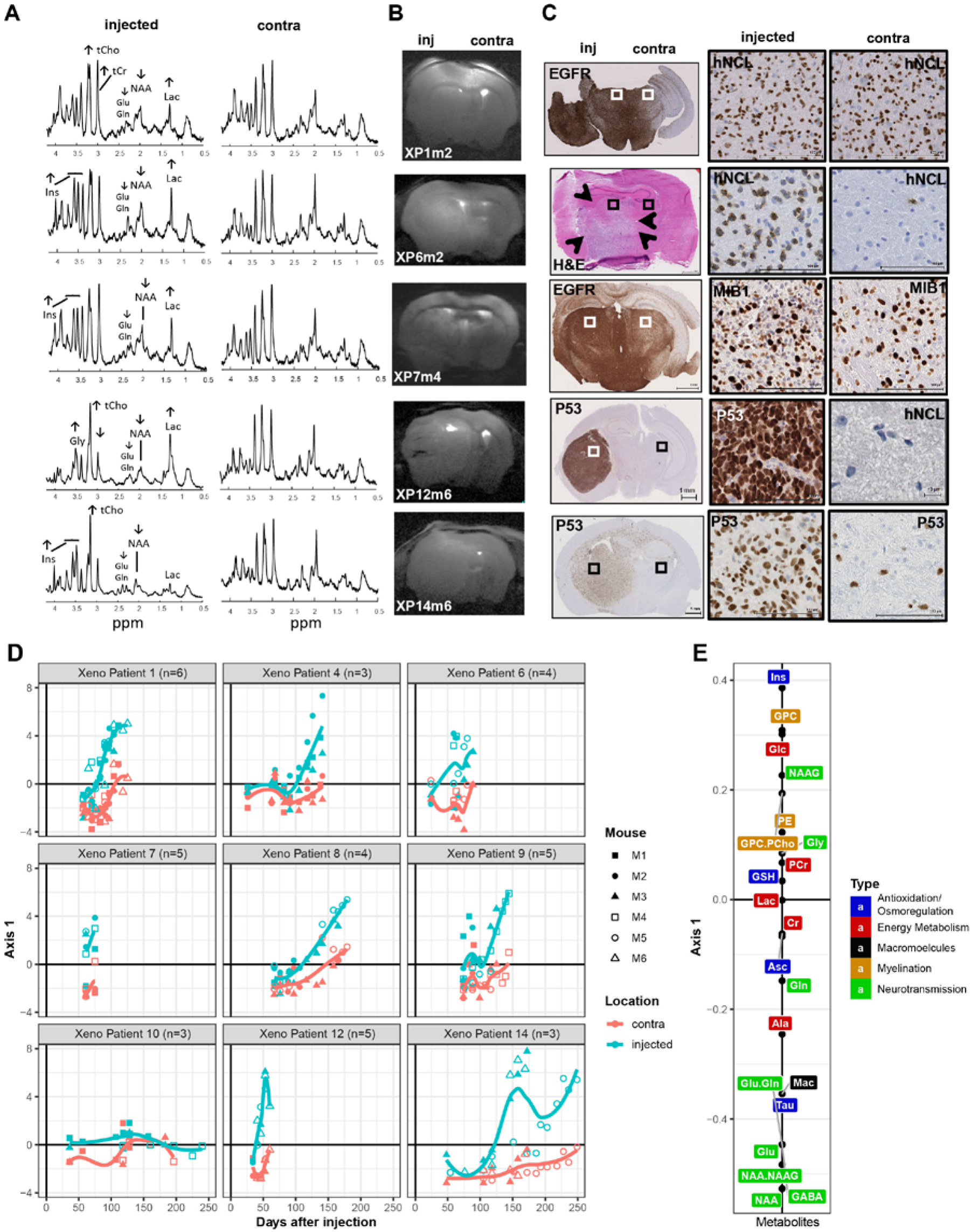
Longitudinal metabolite changes indicate tumor development and invasion. (A) The spectra of the last scans of the injected and the non-injected contralateral side (contra) are displayed, labeled for main metabolites and their changes (indicated by arrows). The corresponding MRIs (B) are annotated for patient origin (P) of the xenografts (X), and mouse number (m), and the indication for the injected (inj) and contralateral side (contra). The histology (C) shows the invasive pattern of the corresponding representative xenografts, with close-up for characteristic areas (squares) of the injected and contralateral side. The human GBM cells are visualized by immunostaining against human specific nucleolin (hNCL), P53, MIB-1, or EGFR. (D) The longitudinal measurements of the metabolite profiles of all mice were analyzed together using STATIS. The metabolite profiles are displayed for all mice by patient against time from injection (days). Each point is a measurement of an individual mouse, identified by a specific symbol as indicated, and corresponds to the respective coordinate of the metabolite profile on the first axis of the STATIS compromise. The 21 metabolites of the profiles are represented on the first axis of STATIS as indicated in panel (E). Metabolite profiles are indicated in blue for the injected, and in pink for the contralateral side for each PDOX-series. The temporal trends are visualized by loess regression. The colors of the metabolites (E) correspond to their major function as indicated (energy metabolism, red; myelination, yellow; macromolecules, black; neurotransmission, green; anti-oxidation & osmoregulation, blue). Abbreviations: N-acetyl aspartate (NAA), N-acetylaspartylglutamate (NAAG), glutamate (Glu), glutamine (Gln), glycerophosphocholine (GPC), phosphocholine (PCho), glucose (Glc), glycine (Gly), myo-inositol (Ins), creatine (Cr), phosphocreatine (PCr), lactate (Lac), glutathione (GSH), taurine (Tau), alanine (Ala), aspartate (Asp), ascorbate (Asc), phosphorylethanolamine (PE), acetate (Ace), gamma aminobutyric acid (GABA), macromolecules (Mac).

The measurements preformed at ultra-short echo-time combined with the increased spectral resolution at 14.1T allowed reporting additional changes of metabolites with low concentrations and overlapping signals that include metabolites like GABA, glutamine (Gln), GPC (dominating the increase seen in tCho), glycine (Gly) in a single measurement without the need of specific editing acquisition sequences (Fig. 1A, E, Supplementary Fig. 1). Of note, these metabolites have been rarely reported in previous studies mainly due to technical limitations.

The profiles of the individual metabolite concentrations, averaged over all PDOX by patient, are displayed in Supplementary Fig. 1. This illustrates the differences in individual metabolite changes, induced by tumor growth in the injected side of the brain and invasion of the contralateral side, between patients, and the similarity among mice injected with tumor cells of the same patients. Absence or minimal increase of lactate (Lac) was observed for xenografts derived of most patients’ tumors, in line with the invasive nature and absence of a necrotic tumor core in the mouse xenografts [21]. The growth pattern of the tumors was patient-dependent, and ranged from diffuse growth with migration across the corpus callosum and invasion of the non-injected side to well delineated tumors with only local invasion (Fig. 1C). The PDOX from P12 yielded the most compact tumors, associated with the highest increase of Lac, tCho, Gly and decrease in Gln, Cr, PCr, Ins among all PDOX (Supplementary Fig. 1), and displayed some Gadolinium uptake in T^1^w-imaging (not shown).

### Spatial metabolite patterns of human GBM resemble temporal evolution of metabolite profiles of PDOX development

To evaluate the metabolite patterns across different areas of the patients’ GBM, MVS was performed at high magnetic fields (7T). For three patients, MVS was acquired sampling an array of 16 to 30 voxels covering distinct regions of the GBM, encompassing parts of the tumor core, the presumed infiltration zone, and adjacent brain (Fig. 2A). This allowed the measurement of 16 metabolites with their corresponding spatial information (Supplementary Table 1), as visualized for nine individual metabolites in distinct heatmaps projected onto the respective MRI of P1 (MRSI, Supplementary Fig. S2). Benefiting from the increases in sensitivity and spectral resolution at 7T together with the short-TE used, we could characterize metabolites beyond those commonly reported at 3T (i.e. NAA, tCho, tCr, Glu+Gln and Ins), including a separate measure of Glu and Gln, and low concentration metabolites such as glycine (an important glioma marker), NAAG, PE and GSH. To determine the spatial pattern of the metabolite profiles, the MVS data of the three patients’ GBM was analyzed simultaneously by STATIS, for which the corresponding representation of the metabolites on the first axis is depicted in Fig. 2B. The metabolite profile attributed to each individual voxel, defined by their coordinates on the first axis of STATIS, was then projected as a heatmap onto the MRIs of the three patients to visualize the spatial organization of the metabolite profiles in the tumor (Fig. 2A).

**Fig. 2.**
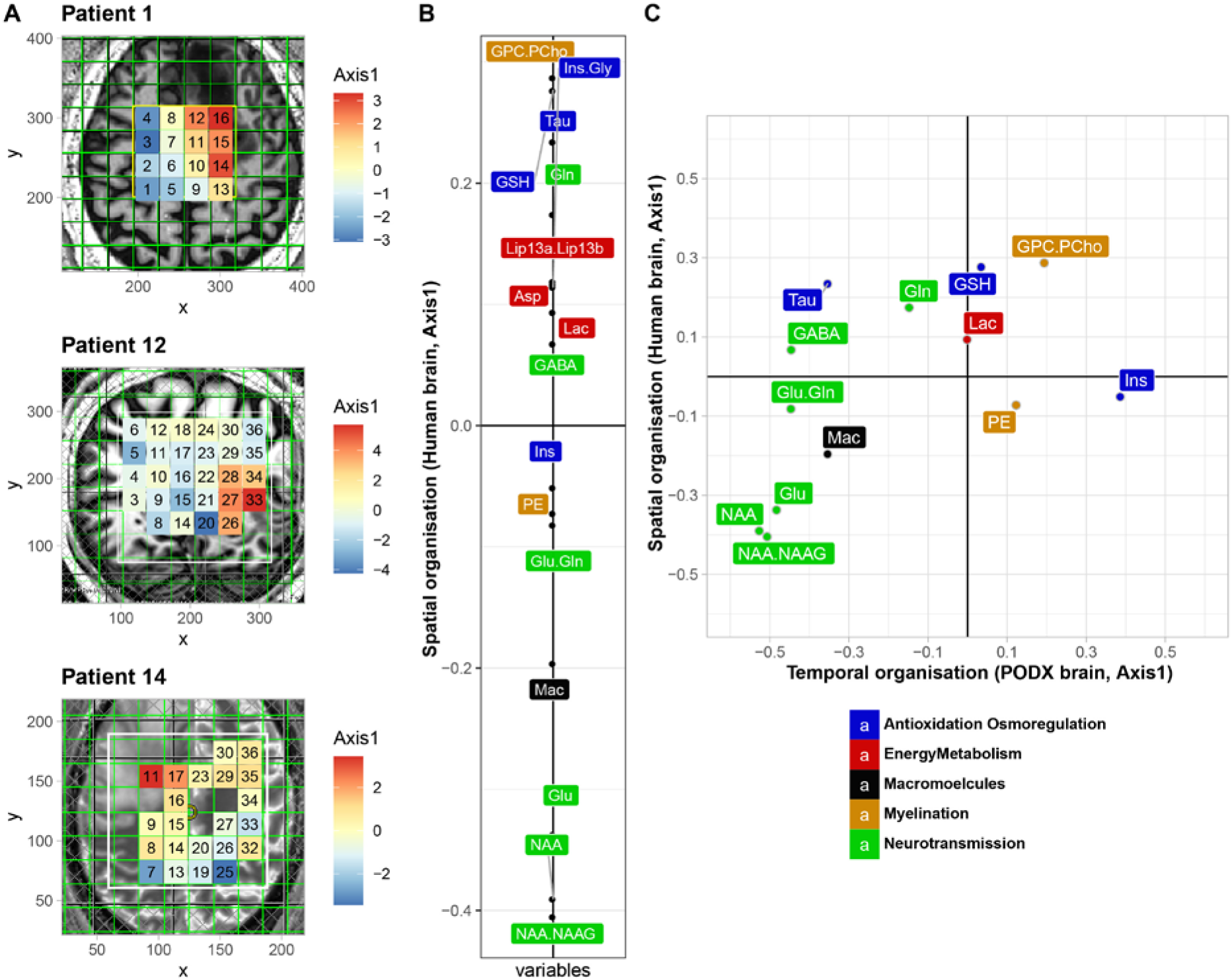
Spatial metabolite profiles determined by multi-voxel analyses. (A) Multi-voxel spectroscopy (MVS) data was acquired for patients P1, P12 and P14. The metabolite spectra acquired were simultaneously analyzed by STATIS, and the respective coordinates from the first axis were then projected as a heatmap onto the corresponding voxel on the respective MRI. The color gradient corresponds to the coordinates projected on the first axis of STATIS, as indicated. The most malignant parts of the tumors are located in the “orange-red” areas. (B) The first axis of STATIS shows the organization of the compromise of the 16 metabolites from MVS analyses of the three patients. The color code of the metabolites corresponds to their major function as indicated. (C) Comparison of the first axis of the 13 common metabolites between human MVS (spatial organization) and mouse metabolite data (temporal organization) is shown in a scatter plot, and displays a remarkable similarity (Spearman correlation = −0.68, p < 0.013). Abbreviations as in Fig 1.

Globally, the spatial pattern of the metabolite profiles was characterized by low levels of NAA, and neurotransmitters, and high concentrations of GPC.PCho, Gln and lipids for voxels located in the tumor core, corresponding to high values on this first axis of STATIS (Fig. 2B) (red in the heatmap, Fig. 2A). In contrast, the probable infiltration zones and “normal” brain were reflected in low values on this first axis (blue in in the heatmap, Fig. 2A) with NAA as a prominent marker (Fig. 2B). To elucidate the resemblance of the spectral patterns of the patients’ GBM, composed of necrotic and infiltrative tumor regions, with the spectra from the PDOXs we analyzed the 13 common metabolites (less than 50% missing values). To this end, the correlation of the coordinates of the common metabolites of the respective STATIS analyses was determined as depicted in a scatter plot (Fig. 2C). This revealed a remarkable similarity (Spearman correlation = 0.68, p < 0.013) between the spatial metabolite profiles derived from MVS across the different areas of the human glioblastoma and the temporal metabolite profiles of the PDOX, reflecting early and late stages of tumor development and invasive growth. Hence, the observed alterations of the metabolite profiles across the sampled (MVS) areas of the patients that encompassed “normal” brain, invasion zone, and the tumor core, resembled the temporal changes of the metabolite profiles in the mouse brains during tumor development, from normal brain, invasive growth, to the full blown orthotopic xenografts at end-stage. This reflected the progression of the metabolite profiles from high NAA/low GPC.PCho (more “normal”, low values on the first axes of human MV and PDOX), to low NAA, GABA, Glu and high GPC.PCho, and Gln (more “tumoral”, high values on the first axes of human MV and PDOX), and remarkably respecting the gradient of the metabolites.

### Gene expression profiles integrating human and mouse reads

To investigate the molecular underpinnings of tumor invasion the human glioblastoma (n=8, excluding P10) and the macro-dissected PDOX of the injected and corresponding region of the contralateral side, corresponding to patients P1, 7, 12, and 14, were subjected to RNA-sequencing (Table 1, Supplementary Table 2). The human and mouse reads, classified using the Disambiguate algorithm [23] were proportional to the presence of human cells in the mouse brain, as estimated on the DNA level with species-specific PCR and the ratio of human tumor cells/mouse cells (Spearman correlation =0.867, p<0.001) as determined by immunohistochemistry with a human-specific nuclear marker (huNCL). Analyzing the human reads only, the genomic variant analysis (SNVs) established the filiation of the original patient tumors and the corresponding PDOX [24]. Characteristic molecular features of the parental tumors, such as mutations or previously described expression signatures [25] (e.g. associated with EGFR overexpression), were mostly retained in the PDOX (Supplementary Fig. 3).

**Fig. 3.**
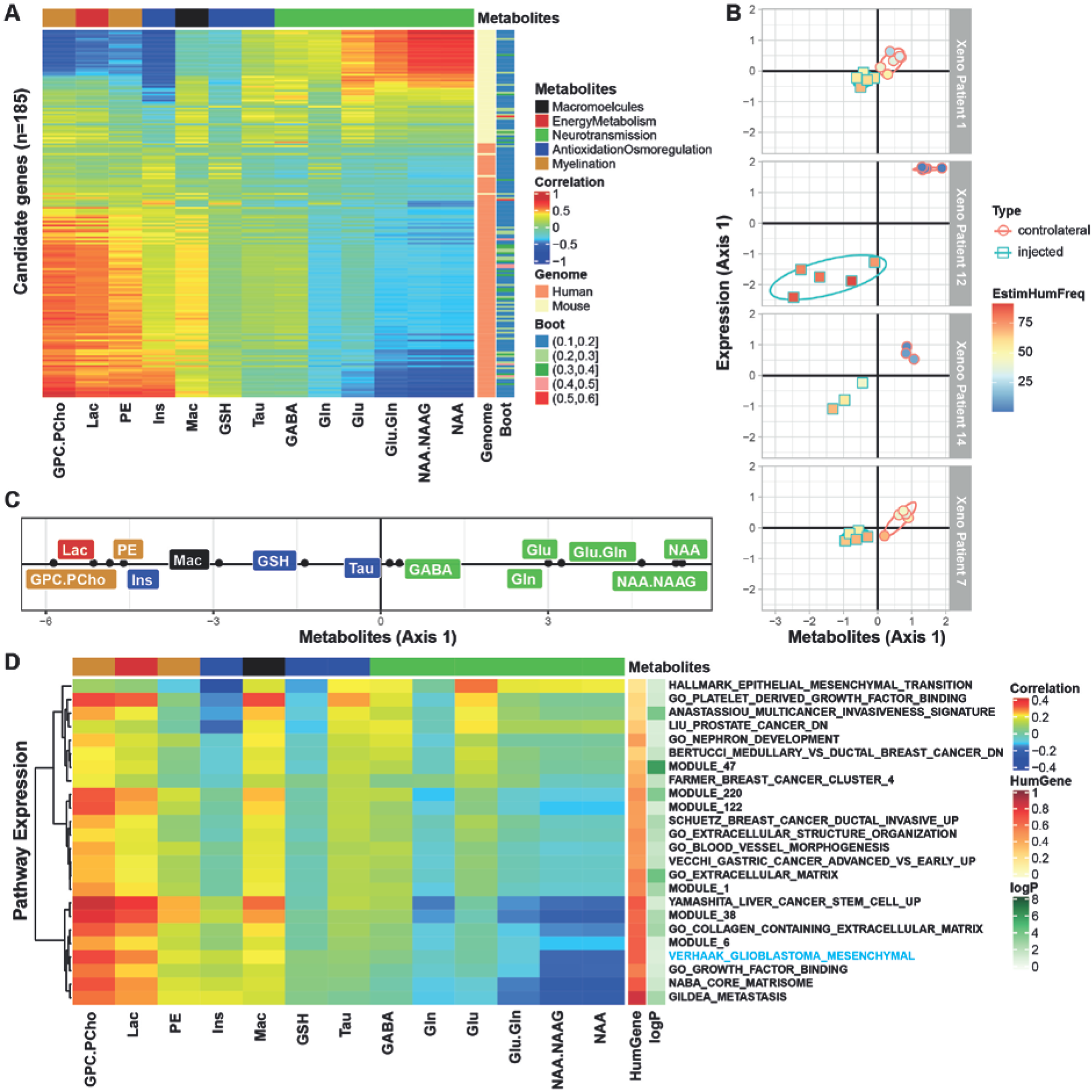
Correlation structure between metabolite profiles (^1^H-MRS) and associated transcriptome (human and mouse). The heatmap (A) illustrates the correlations between 13 metabolites from the last scans of the xenograft bearing mice (injected and contralateral side) and the 185 metabolite associated genes selected by SPLS and retained by the bootstrap procedure (≥0.1). The correlation matrix (coinertia) between genes (rows) and metabolites (columns) is ordered by the 1^st^ axes obtained from coinertia. For each gene the species origin (human or mouse) and the frequency of selection by bootstrap are annotated on the right. (B) The relation between metabolite profiles and gene expression is visualized for all PDOX samples per patient on the vectorial plane defined by the coordinates on the first axes of gene expression and the metabolite profiles from the coinertia analysis, respectively. The samples from the injected side are represented by squares, and by circles for the contralateral side, the color gradient of the symbols indicates the percentage of human reads. (C) The metabolites projected on the first axis of the coinertia analysis. (D) Similarly, the correlation structure between metabolite profiles (^1^H-MRS) and averaged expression of the significant pathways emerging from GSEA (p ≤0.1) using the MSigDB is illustrated in a heatmap (D). The pathways are annotated with the adjusted p-value (p≤0.1), and the proportion of human genes contributing to the pathway.

Integration of mouse and human reads, expectedly revealed that the first axis of the PCA of all samples, based on a gene set selected by sparse PCA (SPCA, 274 genes, selected with lasso regression, consolidated bootstrap; see flowchart of analysis in Supplementary Fig. 4A) was dominated by the species origin, which at the same time reflected the tissue type (tumor vs mouse brain) (Supplementary Fig. 5, Supplementary Table 3). In line, pathway analysis, using gene set enrichment analysis (GSEA) and the molecular signature database (MSigDB), for which the mouse genes were converted into the human homologs, revealed that the top 25 pathways were dominated by cell cycle, proliferation, EGFR signaling, and tumor progression associated gene signatures, and were expectedly mainly described by the human/tumor derived genes (Supplementary Table 4). Interestingly, a tumor invasion related signature was among these top 25 pathways, with similar contributions from both mouse/host-derived and human/tumor-derived genes.

**Fig. 4.**
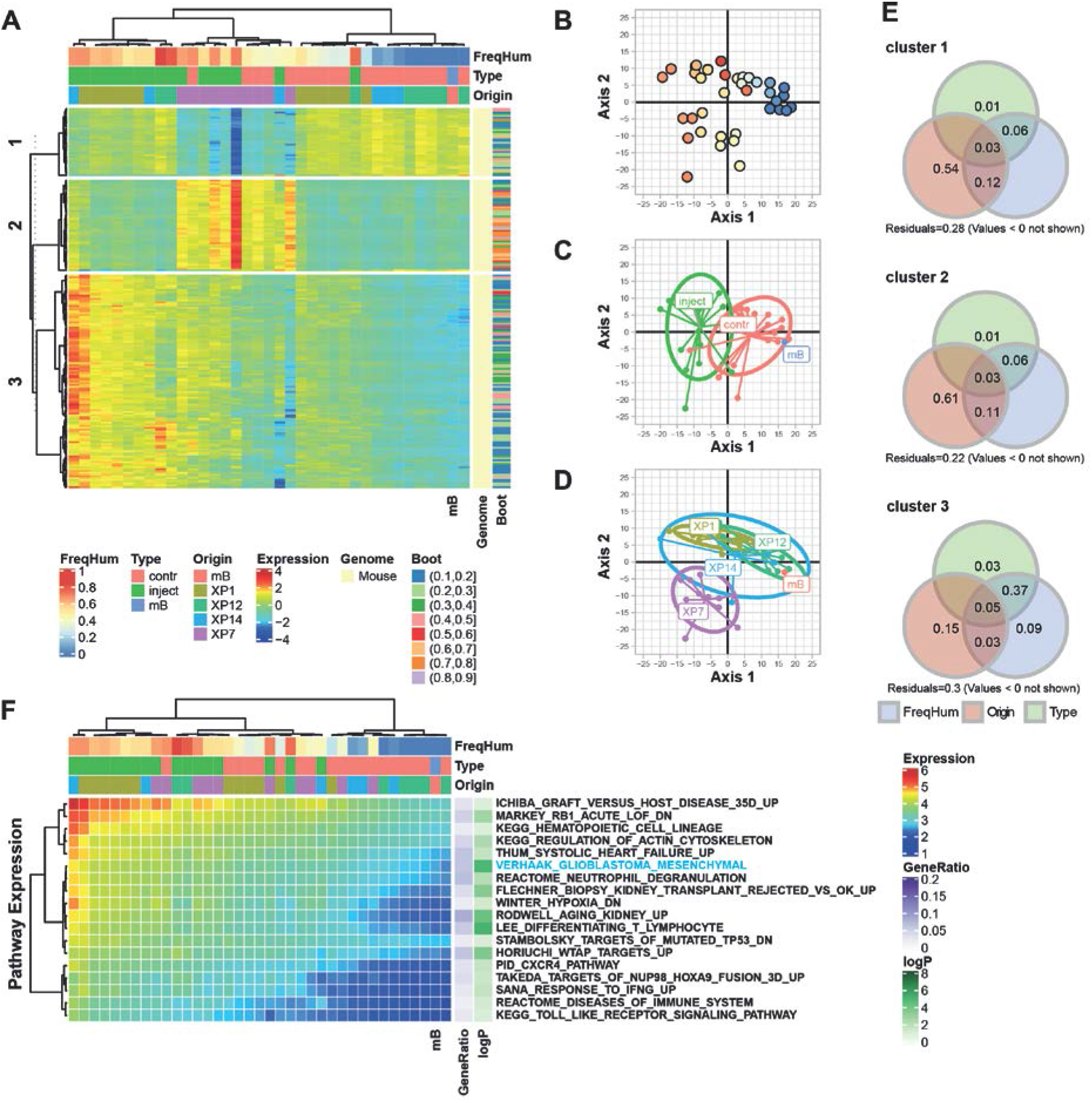
Effect of tumor invasion on the mouse brain transcriptome. Mouse gene expression profiles included PODX samples and one mock injected mouse brain (mB). The heatmap (A) illustrates the normalized expression of the 208 mouse genes selected by SPCA and consolidated by bootstrap (≥0.1). The genes were classified in 3 clusters (consensus k-means clustering for 100 repetitions). (B-D) The samples are projected onto the first two axes of the PCA for the selected genes, where the human read proportion (B), the sample type (C) and tumor origin (D) were added as supplementary variables, annotated with the color code. The contribution of each gene cluster to explain the variance is evaluated by variation partitioning (E) represented by a Venn diagram of variation fractions (percentage) for the three supplementary variables (frequency of human reads, FreqHum; sample origin, Origin; type if tissue, Type). (F) The averaged gene expression of the significant pathways emerging from GSEA (p≤0.1) of cluster 3 is illustrated in a heatmap. The pathways are annotated with the adjusted p-value and the gene ratio (number of selected genes in pathway/number of selected genes in the analysis).

### Molecular pathways associated with changes in metabolite profiles (^1^H-MRS) of invasive tumors

The main interest in this study was to investigate molecular changes underlying tumor development and invasion by integrating gene expression and metabolite information. The latter is amenable to non-invasive analyses allowing longitudinal monitoring of tumor progression and invasion with potential for clinical use.

We determined gene expression profiles associated with the metabolite profiles in the PDOXs acquired from the injected and contralateral side at the last MRI/S scan (end-stage) (see flowchart of analysis in Supplementary Fig. 4C). The patient specific effects were removed in the expression data using within-group PCA (WCA) [26] in order to focus on features of tumor host interaction. A set of 185 genes associated with the metabolite profiles was selected using sparse Partial Least Squares analysis (SPLS) [27, 28], consolidated by bootstrap) that allows combination of different data types linked to the same samples (common column of both tables). The gene set comprised 60 mouse and 125 human genes (Supplementary Table S5). A heatmap illustrates the strong association between the 13 metabolites obtained by MRS and the 185 selected genes in the cross platform comparison (cross table coinertia analysis; coefficient of vectorial correlation, RV 0.73, p=0.01, Fig. 3A). Representing the PDOX samples in function of their metabolite and gene expression profiles (defined by their coordinates on the first axes of the coinertia analysis) revealed a clear gradient following tumor progression, from normal brain (high in NAA) to tumor (low in NAA, and high in choline compounds and enhanced Lac), and change of gene expression (Fig. 3B, C). The extreme features were most prominent for XP12 that yielded the most compact xenografts with some contrast enhancement, while in the contralateral side tumor spread was neither detectable by MRS nor subsequent histology. Of note, the detection of metabolites by MRS is agnostic to the species origin. The RNA coverage revealed a drift from mouse to human reads during tumor development, as expected.

Pathway analysis of the 185 selected genes using the MSigDB database, identified a set of 24 significant pathways (p≤0.1, Bonferroni adjusted). The majority of the pathways were associated with extracellular matrix, extracellular structure, tissue remodeling, adhesion, morphogenesis and remodeling of blood vessels, multi-cancer signature(s) of invasion, metastasis, epithelial mesenchymal transition, and including a mesenchymal glioblastoma signature (Supplementary Table 6). Interestingly, expression of both, mouse and human genes contributed to the selection of all these pathways, ranging from 20% to 80% human genes, depending on the pathway. A heatmap visualizes the correlation of the pathways with the metabolite profiles (Fig. 3D). Interestingly, the gene set linked to epithelial to mesenchymal transition was associated with early stages, while gene sets such as those for matrix remodeling, and mesenchymal glioblastoma were associated with metabolite profiles of more advanced stages of tumor growth.

### Pathway analysis of the invaded brain

In order to determine the molecular effects of glioblastoma invasion on the mouse brain, we specifically analyzed the mouse reads that originate from mouse derived cells enclosed in the macro-dissected xenografts, and the mouse cells of the respective mirrored region from the contralateral side, invaded or not by GBM cells. Two samples with <10^6^ mouse reads were excluded. Both samples were from the injected side of the PDOX, derived from P12 that are highly compact and therefore comprise only few mouse cells (Fig. 1). The heatmap of genes selected by SPCA (n=208, consolidated by bootstrap; supplementary Table 3; see flowchart of analysis, Supplementary Fig. 5B) yielded 3 major gene clusters (Fig. 4A). The organization of the samples by gene expression seemed to be driven by the extent and pattern of the tumor invasion the mouse brains were exposed to. This intriguing observation seems impacted by the patient origin of the injected tumor cells, despite the fact that GBM-derived human reads were excluded from the analysis. A PCA illustrates the samples projected onto the first 2 axes of gene expression (Fig. 4B-D). The first axis explains the difference between the injected and the contralateral side dominated by cluster 3 genes. The gradient of expression of these genes are explained by the extent of invasion, as measured by combination of the percentage of human reads in the samples, and the PDOX type (injected or contralateral side) and explains more than 50% of the variance (combined variation fraction >0.5; Fig. 4E). The pathways associated with gene expression of cluster 3 are related to immune response, regulation of cytokine production, including interferon-α, interleukin 6, and cytokine signaling (e.g. via C-X-C Motif Chemokine Receptor 1, CXCR1), and inflammatory response such as response to interferon-γ (Fig. 4F, Supplementary Table 4). Interestingly, it includes the mesenchymal GBM signature as significant pathway that we also identified as significant, when selecting metabolism associated gene expression signatures, as described above (Fig. 3). Of note, this signature, included in the MSiGDB, corresponds to the signature associated with the expression-based classification of the mesenchymal subtype of GBM [29]. The second axis of the PCA seems to be driven by the impact of PDOX of patient P7 (plus 1 PDOX of P14), with a particular expression pattern captured in cluster 2 that is independent of the extent of tumor invasion. A characteristic feature of these PDOX was rapid growth and/or massive invasion of both hemispheres (Fig. 1). The pathways selected with cluster 2 genes were dominated by ribosomal genes that were highly redundant among the numerous selected pathways. They represented generic protein and RNA related processes, such as regulation of translation, cellular localization of proteins, catabolic processes, and pathways linked to infectious disease.

## Discussion

The invasive capacity of GBM plays a key role in the aggressiveness of the tumor, its resistance to treatment, recurrence and poor prognosis. The invasive component is typically shielded by the BBB, and it remains a challenge to monitor tumor infiltration of the brain parenchyma using standard MRI techniques [30, 31]. It is therefore of crucial importance to develop adequate tumor models featuring this invasive part that is considered highly relevant for tumor recurrence.

We evaluated molecular features of invasive PDOX that may serve as proxy for studying the invasive front of GBM. To this aim, we present the first *in vivo* comparison of human tumors and respective PDOX in the mouse brain on the levels of radiological behavior and metabolism, as determined by ultra-high field ^1^H-MRS of the patients (7T) and the respective PDOX (14.1T). We found a good concordance between the temporal changes of the metabolite profiles observed during invasive growth of the PDOX, and the metabolite profiles reflecting the spatial organization of MVS originating from different parts of the human tumors, comprising normal brain, infiltration zone, and the tumor core (Fig. 2C). Hence, metabolic changes may inform on alterations in the infiltration zone that is not visible on routine MRI evaluation. Importantly, highly resolved ^1^H-MRS was more sensitive to detect early development of the highly invasive PDOX than conventional structural MRI acquisitions. This may open the possibility to test and non-invasively monitor early treatment effects in the invasion zone shielded by an intact BBB. Interrogating the molecular mechanisms related to tumor invasion by means of multi-dimensional analysis allowed simultaneous exploration of metabolic and transcriptomic changes of the samples and respective associated pathways. This combined analysis provided insights into the biological mechanisms that are underlying the metabolic changes that can be followed non-invasively. Most importantly, both tumor and host contributed to gene expression profiles associated with the biological pathways uncovered, supportive of their biological relevance. These signatures indicated active processes associated with changes of the extracellular matrix, tissue and blood vessel remodeling, along with signatures attributed to tumor invasion, metastasis, and mesenchymal GBM. Lately, the tumor matrix, and the associated cell-matrix interaction have received more attention in solid extra cerebral tumors [32], while little remains known in brain tumors [33]. Interestingly, gene expression annotated for hallmarks of epithelial/mesenchymal transition was associated with more normal brain-like metabolic features, while the gene expression related to matrisome, metastasis, and mesenchymal GBM were more strongly correlated with metabolite profiles of more advanced tumors (Fig. 3D).

Taking a different view, and evaluating the transcriptome originating from the invaded brain only (mouse reads, Fig. 4), revealed inflammatory signatures and profiles of cytokine mediated regulation of immune response. The mesenchymal GBM signature that was again associated with a more aggressive extent of tumor invasion/tumor burden as estimated by the proportion of human reads, despite the fact that they were excluded from the analysis. The mesenchymal GBM subtype has been associated with more prominent recruitment of macrophages [34] for which the functional interaction with the GBM cells has been described recently, evoking that this interaction drives the mesenchymal-like state of GBM [35]. Along the same lines, testing our previously reported GBM gene expression signatures [25] with GSEA, we found two matching clusters that were significantly associated with the expression profiles of the invaded brain: an interferon-induced gene signature (G12; adjusted p-value, 0.0002), and an innate immune response signature (G24; adjusted p-value, 0.0001). This suggests some concordance of the brain reacting to tumor invasion in the present PDOX model and human GBM.

These molecular insights suggest that the tumor/host interaction of tumor invasion can be modeled by monitoring metabolic changes in PDOX and that these changes present a good match with their human counterparts as described in this study. The insights derived from temporal metabolic changes associated with diffuse tumor invasion and progression may be applicable to help monitor the invasion zone in patients to identify early responses to treatment or on the contrary, early tumor progression.

The obvious limitations are the lack of an intact immune system in this mouse model that is obviously expected to play an important role in tumor progression and treatment [36].

The PDOX may represent an attractive perspective to develop and evaluate drugs aimed at treating tumor cells in the invasion zone. Other invasive orthotopic brain tumor models may also be suitable for metabolic monitoring. For instance, we have reported similar longitudinal metabolic changes using established GBM derived sphere (GS) lines yielding highly invasive orthotopic xenografts [21]. The changes in the metabolic profiles were sensitive enough to follow the distinct evolution when transducing the GS cells with a tumor suppressor gene (Wnt inhibitory factor 1) [21]. This is important, as the median latency of freshly resected patient derived tumor cells to fully develop into PDOX is generally too long (5-35 weeks in this study) to serve as faithful avatars for patient specific testing of targeted drugs to guide treatment choices.

Taken together, invasive PDOX models exert spectroscopic and transcriptomic features of brain infiltration that show similarities with the presumed infiltration zone of GBM. This supports their suitability as relevant models for studying the non-enhancing part of GBM. The possibility of non-invasive *in vivo* monitoring of invasive growth in this difficult to evaluate compartment, may allow early detection of relapse and monitoring of treatment effects of novel drugs that eventually may be translated into the human setting.

## METHODS

### Patient selection and ^1^H MRS of patients

Patients planned for surgery of a suspected GBM at the Lausanne University Hospital (CHUV) were enrolled (clinicaltrials.gov, NCT02904525) with written informed consent. The study was conducted in accordance with the Declaration of Helsinki, and the protocol was approved by the local ethics committee CER-VD (F-25/99, 268/14).

Patients underwent ^1^H-MRS/I in a 7 Tesla/68 cm MR-scanner (Siemens Medical Solutions, Erlangen, Germany). B0 field homogeneity was optimized using FASTMAP [37]. A 32-channel receive coil (NOVA Medical Inc., MA) with a single channel volume transmit coil or a ^1^H two loops surface coil were used depending on the location of the gliomas. 3D T1-weighted MR images acquired using MP2RAGE (TE/TR = 3.37/5000 ms, TI1/TI2 = 700/2200 ms, slice thickness = 1 mm, FOV = 176 × 256 mm2, matrix size = 176 × 256) [38] were used to position the volume of interest (VOI) for MRS measurements. Single voxel ^1^H MR spectra (SVS) were obtained using the semi-adiabatic SPECIAL localization sequence [18, 39] with TE = 16ms, TR = 5.5−8.5s (depends on the SAR restriction), and number of averages = 48-96. 2D multi-voxel ^1^H MRSI was measured by a sEmi-Adiabatic Spin-Echo MRSI sequence (EASE) [40] for P1, P12 and P14, using the following parameters: TE=16ms, TR=4.3s, NA=1, FOV=200×200mm^2^, VOI= 60×60mm^2^, slice thickness=15mm, matrix=16×16, elliptical k-space sampling. Water and lipid suppression techniques were applied prior to the localization using VAPOR with OVS according to consensus recommendations [41].

^1^H MRS spectra were analyzed by LCModel [42] using a basis set with simulated metabolite spectra and an experimentally measured macromolecule baseline [43]. LCModel simulated macromolecule and lipid components were used during the analysis to allow fitting for potential lipid and MM resonances that arose from tumors. Due to time restriction in patient scans, additional water acquisition for normalization purposes could not always be performed. Metabolite ratios to total creatine were calculated for both SVS and MRSI measurement. Metabolites that were quantified with CRLB > 50% were excluded in the analysis.

### Orthotopic Mouse Glioma Model

Tumor tissue obtained at surgery was split in two, one part was frozen for subsequent analyses and the second part was dissociated into single cells and re-suspended in stem cell media (DMEM/F12 supplemented with B27 and growth factors) as described [44]. The next day 10^5^ cells were transplanted stereotactically in a volume of 5 μl (Hanks’ balanced salt solution, HBBS, with phenol red, no calcium, no magnesium; Thermo Fisher Scientific) into the striatum (coordinates: bregma 0.5mm anterior, 2mm lateral and 3mm ventral)[45] of male immunocompromised mice (n=3-6/patient; age, 6-8 weeks; NOD-SCID gamma knock-out mice, NOD.Cg-Prkdcscid II2rgtm1Wjl/SzJ, bred in-house) using a micro pump (injection rate 5 μl/min, Stoelting). For the procedure, anesthetized mice were placed into a stereotactic frame, and fixed with a mouth piece (Stoelting) as previously described [46]. Mice were checked daily and sacrificed at first signs of neurologic symptoms (lethargy, ataxia and seizures) or body weight loss (>10%). All animal procedures were performed under anesthesia/ analgesia, and protocols were approved by the concerned Swiss authorities (VD-1181_6; VD-2777).

### *In vivo* ^1^H-MRS of orthotopic xenografts in the mouse

^1^H-MRS experiments were carried out in a 14.1 Tesla animal scanner with a 28-cm horizontal bore (Agilent Technologies, Palo Alto, CA, USA) using the SPECIAL sequence (TR = 4s, TE = 2.8ms, 160 or 240 scans) [18] as previously described [21]. An axial T2-weighted image (fast spin-echo sequence, TR/TE = 5000/13 ms, FOV =18×18 mm, slice thickness 0.6 mm, 6 averages) was acquired before the ^1^H-MRS for voxel positioning (VOI, 2×2×2 mm^3^), centered in the striatum of the injected, and symmetrically, in the contralateral hemisphere. SVS was preferred over MVS due to the location of the tumors and the shorter acquisition time, thus allowing repeated measurement in the same animal. Modifications in the BBB were assessed in selected cases with T1-weighted coronal fast spin-echo images with Gadolinium contrast (Gadovist 1.0, Bayer, Leverkusen, Germany).

The first scan was performed 6 weeks after injection or at onset of symptoms (earliest week 4), and was repeated every 1-2 weeks. Spectra were quantified using the LCModel [21, 42] using a simulated basis set of brain metabolites combined with an experimental spectrum of macromolecules (Mac) acquired in a healthy subject and simulated lipids (at 0.9, 1.3 and 2.0ppm) when present according expert recommendations [47]. Data shown are selected for accurate quantification, following the criterion Cramer-Rao lower bound (CRLB) < 40%. Metabolites were normalized to tCr unless stated otherwise. Metabolite concentrations using water as internal reference were also computed and used in Supplementary Fig. 1.

### Tissue Processing and Immunohistochemistry

At final sacrifice, the fresh mouse brain was cut using a brain mold. The central coronal brain slice (~5mm) encompassing the injection-site was frozen in OCT (Tissue Tek), and the rest was formalin fixed and paraffin embedded for histology. Serial frozen sections were cut on a cryostat from the central slice of the mouse brains. The first and last sections were used to determine tumor location and estimate tumor cell content, based on hematoxylin and eosin (H&E) staining and nuclear immune-positivity for human-specific NCL (ab13541, Abcam, Cambridge, UK) as previously described [21]. Xenografts were macro-dissected, guided by H&E and NCL staining, and collected from the injected and contralateral side separately. In absence of apparent spread to the contralateral side, the tissue of the symmetric area on the contralateral side was collected. Immunohistochemistry was performed for EGFR (E30), Ki67 (MIB1), GFAP (G-A-5), and TP53 (DO-7) (platform Ventana, Roche).

### DNA/RNA isolation, PCR, and RNA sequencing

RNA/DNA was isolated from the macro-dissected xenografts (injected/contralateral side, separately) and human GBM (AllPrep DNA/RNA Mini Kit, Qiagen). The ratio of human/mouse cells in the xenografts was estimated by species specific PCR (DNA) [48]. Library preparation and RNA-sequencing was performed at the Lausanne Genomic Technologies Facility (LGTF, University of Lausanne; TruSeq Stranded Total RNA Library Kit; Illumina HiSeq 2500). The samples were barcoded and randomized between lanes for sequencing. For samples with a tumor/mouse cell ratio of ~50%, we aimed at 60×10^6^ reads, and for all other samples, including the human GBM 30×10^6^ reads.

### Data preparation and analysis of RNA sequencing

Preprocessing of RNAseq data was performed following the standard pipeline and recommendations from bcbio-nextgen (version 1.0.4, http://bcbio-nextgen.readthedocs.org/en/latest/). Reads were aligned to the human (GRCh37) and mouse reference genome assembly (mm10) by hisat2 aligner (version 2.1.0), and classified into three classes (mouse, human, ambiguous) by the Disambiguate algorithm [23]. Transcripts with low read counts, or classified as ambiguous were removed. The gene expression data were summarized by trimmed means of M-values (TMM) of normalized counts (R package edgeR) [49, 50], including log-transformation and using read counts and full library size. The Variant calling analysis was performed with the software VarDict [24] and the genomic variant annotations were obtained by SNPeff [51]. SNVs with low quality estimation and silent and synonymous mutations were excluded. The genes listed in the COSMIC database as mutated in glioma and glioblastoma [52] were used for our analysis. In addition, the SNVs identified were compared with the mutation database from TCGA for Glioma and Glioblastoma (R package RTCGA.mutations). For personal privacy reasons the RNA-sequencing raw data will be made available upon request.

### Data analyses and variable selection

The metabolite profiles and expression data were examined by principal component analysis (PCA) and heatmap representation based on Euclidean distance and Ward’s algorithm for clustering. Missing values were imputed by regularized iterative PCA algorithm [53]. The global differences between groups were tested by a Monte-Carlo test (permutation) on the between-groups inertia percentage [54].

The MVS in patients and longitudinal metabolic profiles in mice were analyzed by STATIS [55, 56] that allows simultaneous analysis of different data arrays, matched by common columns (same variables) based on principal component analysis (PCA). Briefly, in using inertia operators and RV-coefficients [52], STATIS compares the “images” (interstructure) of each dataset, to find a consensus (compromise) and simultaneous representations of each dataset on the same factorial plane (intrastructure).

A flowchart details the strategies for gene selection and procedures used for data integration with metabolites and gene ontology (Supplementary Fig. 4). Gene expression was analyzed by sparse PCA (SPCA) [57] for gene selection, using singular value decomposition and lasso regularization (Supplementary Fig. 4A-B). The gene signature was consolidated by bootstrap (50 repetitions). At each iteration, two components were retained to describe data organization, and 50 variables (genes) were kept in each loading vector [57]. Thus, the most frequently selected genes were retained (cut-off **≥**0.1). The correlation structure between metabolite profiles and gene expression, was investigated by sparse Partial Least Squares analysis (SPLS) [27, 28] after removing unwanted effects of tumor origin by within-group PCA (WCA) [26] (Supplementary Fig. 4C). The SPLS approach combines both integration and additional variable selection (lasso regularization) simultaneously on two data sets in a one-step strategy. The selected gene-set was consolidated by bootstrap (50 repetitions). At each iteration, the association between metabolite profiles and gene expression was summarized by two components, for which all metabolites were retained and 50 variables (genes) were kept in each loading vector. Finally, the most frequently selected genes were retained (cut-off **≥**0.1). The Coinertia analysis [58], a multivariate method for coupling two tables, summarizes the correlation structure between metabolite profiles (^1^H-MRS) and expression of the selected genes (R packages ade4 and mixOmics). The common correlation structures between gene expression and metabolite profiles were investigated by permutation test and reported as RV-coefficient (vectorial correlation coefficient) [59].

Gene set enrichment analysis (GSEA) was performed with the molecular signature database (MSigDB v7.0, updated August 2019, all 8 collections) [60] using hypergeometric tests (R packages msigdbr and ClusterProfiler). Gene-sets with Bonferroni adjusted P-values≤0.1 were considered significant. The conversion of mouse genes into the corresponding human homologs was performed with R package biomaRt [61]. All analyses and graphical representation were performed with R version 3.6.1 (URL http://www.R-project.org) [62] and the R packages survival [63], missMDA and ade4 [64].

## Funding

The study was funded by the Swiss Bridge Award, the Swiss Cancer Ligue (KFS-3998-08-2016, KFS-4461-02-2018), the Swiss National Science Foundation (SNF-3103-163297, SNF-3103-182821), Harry’s ride and Cadot foundations.

## Conflict of interest statement

The authors declare to have no competing interests.

## Authorship statement

C.C., A.F.H, and M.E.H conceived and designed the study; P.B. designed, performed the biostatistical analyses, and interpretation; L.X. performed MRI/MRS (7T) of patients; C.C. and M.La. performed MRI/MRS of mice; O.G., and M.-F.H. performed transplantations, and sample analyses; M.Le. assisted animal work; R.T.D. provided surgical specimen and J.P.B. neuropathology expertise; A.F.H. lead the patient study; M.E.H. coordinated data integration, wrote the manuscript with P.B.. All authors contributed to data interpretation and manuscript writing.

## Acknowledgements

We thank the patients and their families for their support and participation. Information about TCGA can be found at “http://cancergenome.nih.gov”. We acknowledge access to the facilities and expertise of the SIB, and the CIBM.

## Notes

### Competing Interest Statement

The authors have declared no competing interest.

