## Supplemental Material for "Metabolic and transcriptomic profiles of glioblastoma invasion revealed by comparisons between patients and corresponding orthotopic xenografts in mice"

*Equal contribution
#shared senior authorship

**Corresponding author**:

### Supplementary Tables (uploaded as separate excel files)

**SupplementaryTable1_Metabo.xlsx :**

- Metabolite datasets

**SupplementaryTable2_RNAseqDescription.xlsx** :

- Description samples (NB reads, categories, etc.)

**SupplementaryTable3_RNAseq.xlsx** :

- List of candidate genes (n=227) for aggregated human and mouse RNAseq data;
- Candidate genes (n=208) mouse only

**SupplementaryTable4_RNAGSEA.xlsx**:

- Pathways associated with gene expression using all samples
- Pathways associated with gene expression restricted to mouse sequences

**SupplementaryTable5_RNAmetabo.xlsx**

- Candidate genes (n=185) associated with metabolites

**SupplementaryTable6_RNAmetaboGSEA.xlsx**

- Pathways linked with metabolite associated gene expression

### Supplementary Figures 1-5

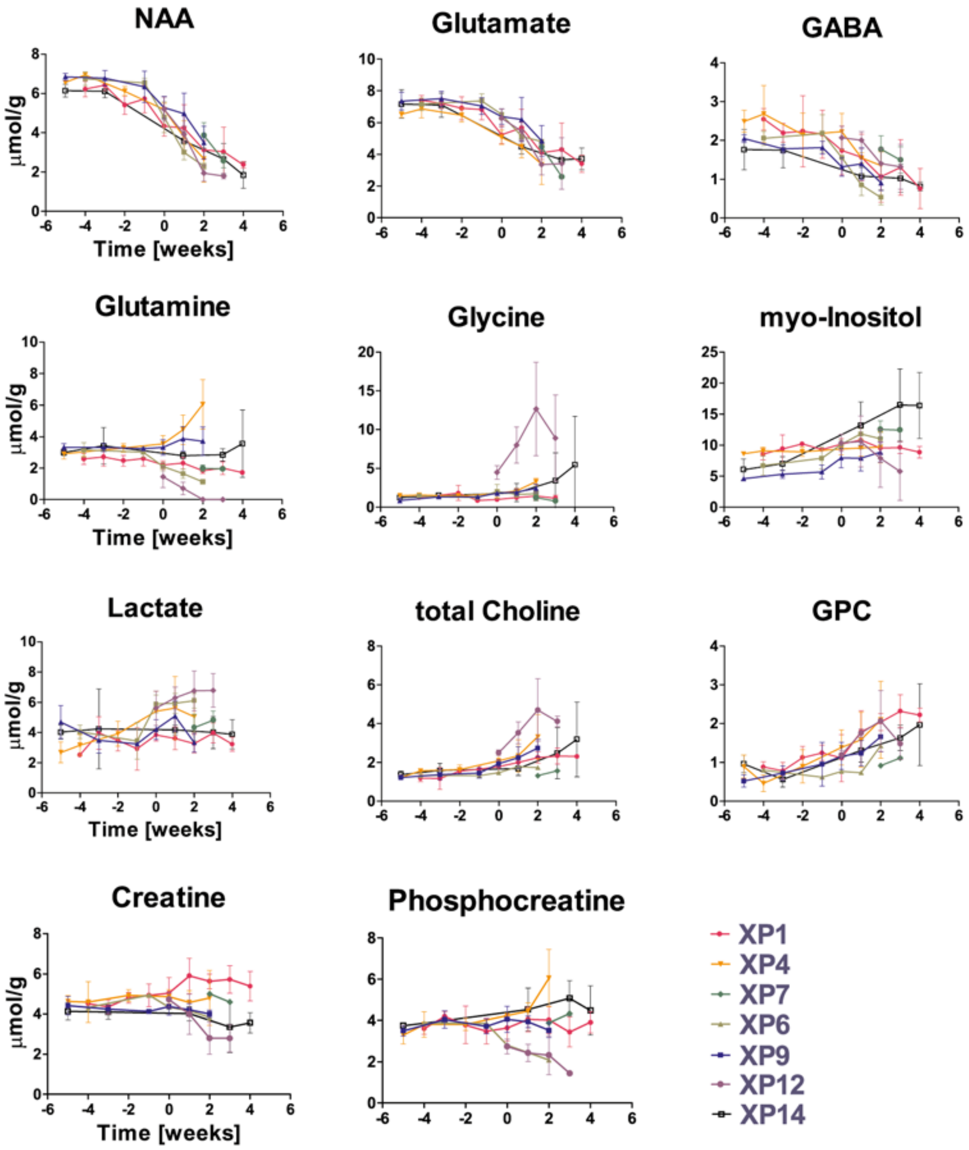
NAA
Glutamate
GABA
Glutamine
Glycine
myo-Inositol
Lactate
total Choline
GPC
Creatine
Phosphocreatine
μmol/g
Time [weeks]
XP1
XP4
XP7
XP6
XP9
XP12
XP14

A

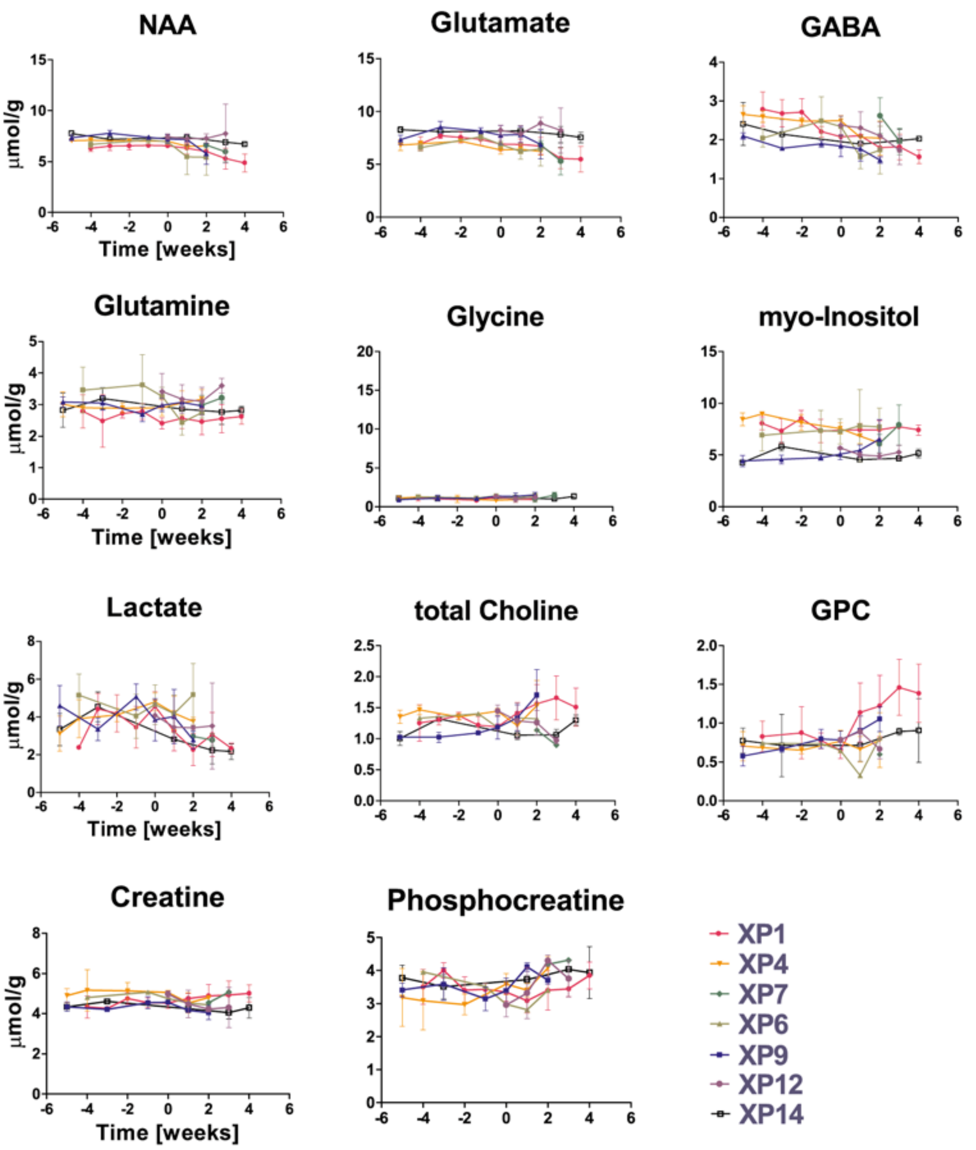


**B**

**Supplementary Fig. 1.** Longitudinal variations of metabolite concentrations of PDOX: (A) injected side, and (B) contralateral side. Points represent the group average by patient at a given time point, error bars are standard deviations. Metabolite concentrations were normalized to internal water. Time is expressed in weeks relative to *first tumor appearance* corresponding to "week 0" - defined as a significant change in NAA concentration on the injected side compared to the first neurochemical profile measured after tumor cell transplantation (Student t-Test on group averages, $p<0.05$). In general tumors cannot be visualized by conventional MRI at this time point. Longitudinal data were registered for all groups except P8 (N=5), where mice M1 and M3 died prematurely during anesthesia, and mouse M4 developed an intracerebral oedema that impeded a correct acquisition of spectroscopic data.

For comparison, the scales of the metabolite concentrations and time points are the same for the injected (A) and contralateral side (B)

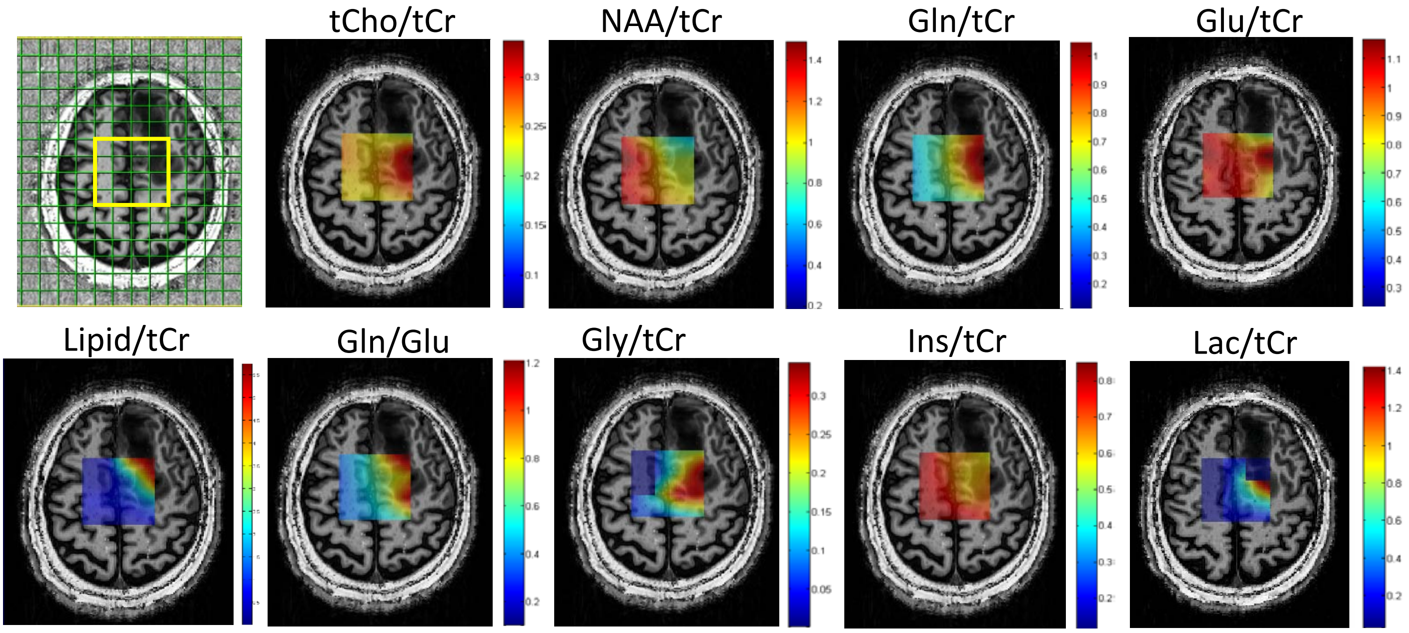


**Supplementary Fig. 2.** Multivoxel $^{1}$H MRS at 7 Tesla of patient 1. Metabolic maps of relative concentrations to tCr for patient P1 visualized in a heatmap projected onto the respective voxels (TE/TR = 16/4290ms, NA = 1, FOV = 200×200mm$^2$, VOI = 60×60mm$^2$, tk = 15mm, matrix = 16×16, elliptical k-space sampling, acquisition time =11min) overlaid with the transverse slice of $T_1$-weighted image (MP2RAGE: TE/TR = 3.37/5000 ms, TI1/TI2 = 700/2200 ms, slice thickness = 1 mm, FOV = 176 × 256 mm$^2$, matrix size = 176 × 256).

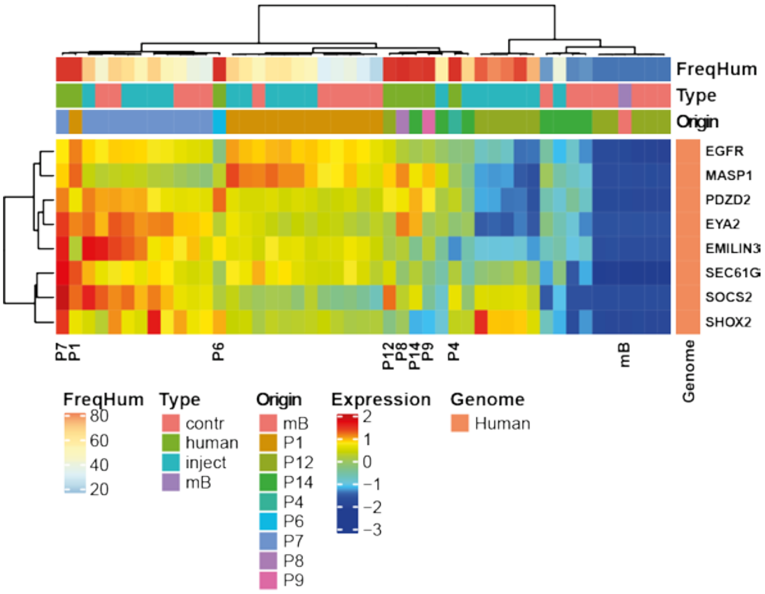


**Supplementary Fig. 3. Retention of characteristic molecular features in PDOX.**

Characteristic molecular features of GBM were evaluated in the patients' tumors and corresponding PDOX. The heatmap visualizes a GBM-derived expression signature associated with *EGFR* overexpression (G25[1]), a characteristic feature of ~50% of GBM. The *EGFR* high signature was retained in the PDOX from two patients (P1, P7), while it was lost/absent in all PDOX of patient P12, which was confirmed on the protein level by immunohistochemistry for EGFR (Hirsch score 0; Table 1). Of note, GBM of P12, expressed a *TP53* mutant (c.743G>A, p.Arg248Gln), a rare combination, as *TP53* mutations and *EGFR* amplifications are usually mutually exclusive [2]. This *TP53* mutation was retained in all corresponding xenografts, in accordance with strong nuclear p53 immunostaining (Figure1). The search for other GBM relevant mutations further revealed a *PTEN* frameshift mutation (10:89720804_ACT/CC) identified in the original tumor and all xenografts of P1, while a *PTEN* mutant (c.388C>T, p.Arg130Ter) was found expressed only in the xenografts, but not the original tumor of P14, suggesting a sub-clonal origin. FreqHum: frequency of human gene expression; mB: normal mouse brain.

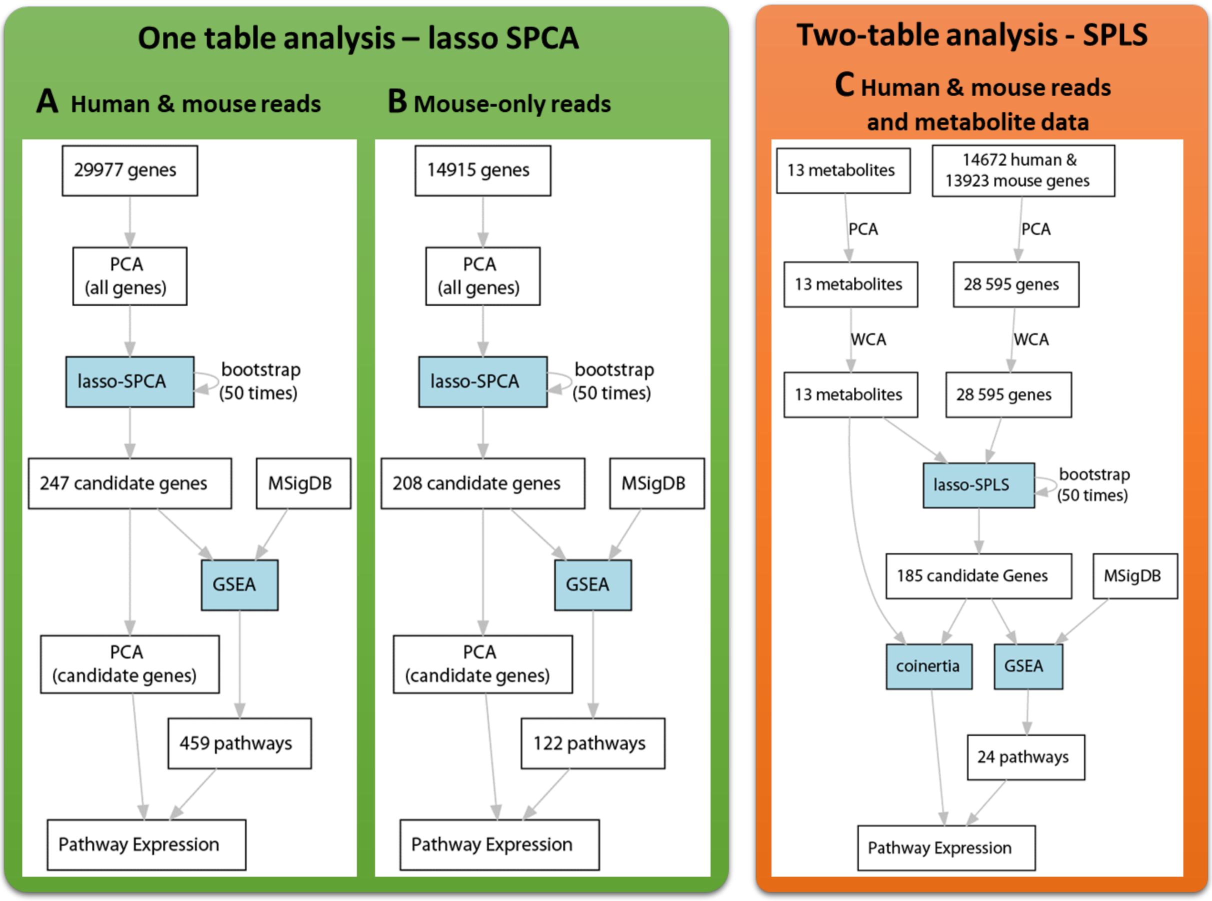


**Supplementary Fig. 4.** Flow charts for gene selection procedures. Description of the two procedures used to detect candidate genes: The candidate genes were selected by Sparse principal components analysis (SPCA) and bootstrap procedure. Candidate gene expression was summarized by principal components analysis (PCA) (**A, B**). ( **C**) The correlation structure between metabolite profiles and gene expression (two-table analysis) from PODX, was investigated by sparse Partial Least Squares analysis (SPLS) [3, 4] after removing unwanted effects of tumor origin by within principal components analysis (WCA) [5]. The SPLS involves the maximization of covariance between two datasets (e.g. transcriptomics and metabolites). Both types of analyses, SPCA and SPLS, include variable selection provided by lasso regularization. The gene signatures were consolidated by bootstrap (50 repetitions) and most frequently selected genes were retained as candidates, with a cut-off superior or equal to 0.1. The molecular signatures database (MSigDB) [6] and hypergeometric tests were used to perform gene set enrichment analysis (GSEA) for the candidate genes. The Coinertia analysis (Coinertia) [7], a multivariate method for coupling two tables, summarizes the correlation structure between metabolite profiles and expression of the candidate genes. The multivariate analyses were performed with R packages ade4 (PCA, WCA and Coinertia) and mixOmics (SPCA1, SPLS).

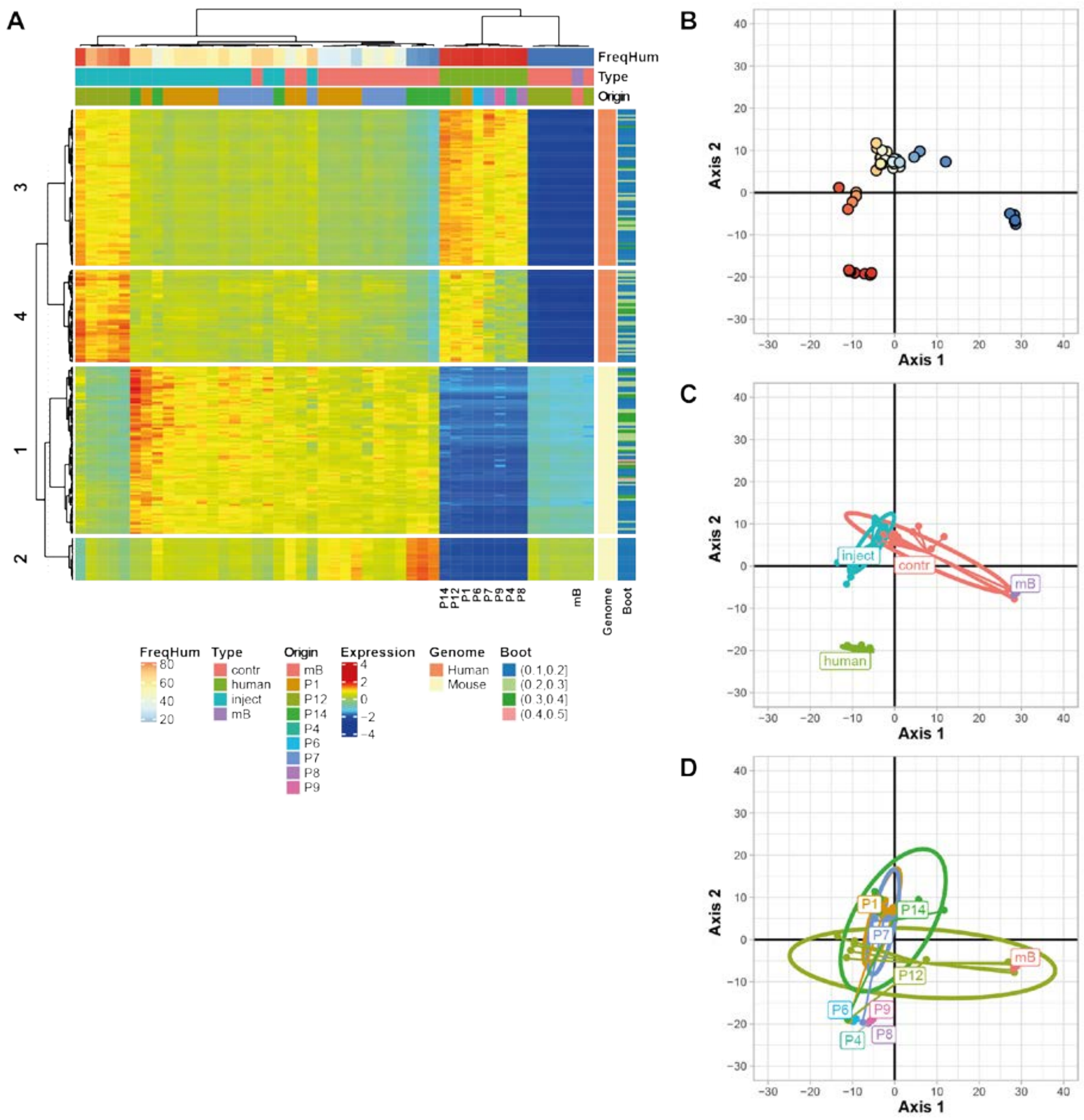


**Supplementary Fig. 5.** Expression analyses including all PDOX and original human GBMs. The heatmap (A) illustrates the normalized expression of the 247 genes (134 human genes and 113 mouse genes) selected by Sparse Principal Component Analysis SPCA based on the combination of log-normalized data of human and mouse reads (counts per million, CPM) and the bootstrap procedure. The samples were ordinated on the two first axes of the sparse principal components (SPCA) where the proportion of human reads (B), the type (injected vs contralateral side) (C), and the tumor origin (D) were added as supplementary variables. FreqHum: frequency of human gene expression; mB: normal mouse brain; Px: patient number
